# Sickle cell disease patient plasma sensitizes iPSC-derived sensory neurons from sickle cell disease patients

**DOI:** 10.1101/2023.01.10.523446

**Authors:** Reilly L. Allison, Anthony Burand, Damaris Nieves Torres, Amanda M. Brandow, Cheryl L. Stucky, Allison D. Ebert

**Author notes:** Corresponding author: Name: Allison Ebert, PhD, Address: 8701 Watertown Plank Rd, Milwaukee, WI 53226, (p).

## Abstract

Individuals living with sickle cell disease (SCD) experience severe recurrent acute and chronic pain. In order to develop novel therapies, it is necessary to better understand the neurobiological mechanisms underlying SCD pain. There are many barriers to gaining mechanistic insight into pathogenic SCD pain processes, such as differential gene expression and function of sensory neurons between humans and mice with SCD, as well as the limited availability of patient samples. These can be overcome by utilizing SCD patient-derived induced pluripotent stem cells (iPSCs) differentiated into sensory neurons (SCD iSNs). Here, we characterize the key gene expression and function of SCD iSNs to establish a model for higher-throughput investigation of intrinsic and extrinsic factors that may contribute to increased SCD patient pain. Importantly, identified roles for C-C Motif Chemokine Ligand 2 (CCL2) and endothelin 1 (ET1) in SCD pain can be recapitulated in SCD iSNs. Further, we find that plasma taken from SCD patients during acute pain increases SCD iSN calcium response to the nociceptive stimulus capsaicin compared to those treated with paired SCD patient plasma at baseline or healthy control plasma samples. Together, these data provide the framework necessary to utilize iSNs as a powerful tool to investigate the neurobiology of SCD and identify potential intrinsic mechanisms of SCD pain which may extend beyond a blood-based pathology.

**Key Points:**

- Sickle cell disease (SCD) stem cell derived sensory neurons (iSNs) recapitulate important SCD phenotypes *in vitro*.
- SCD patient plasma sensitizes SCD iSNs to TRPV1 stimulation by capsaicin.

## Introduction

Sickle cell disease (SCD) is the most common inherited hemoglobinopathy in the United States, affecting over 100,000 individuals who are primarily Black and Hispanic Americans^1–3^, and over 3 million individuals world-wide^1–3^. The underlying pathophysiology of SCD is complex and includes, but is not limited to, red blood cell (RBC) sickling, chronic hemolysis, vaso-occlusion with resultant ischemia-reperfusion injury, and chronic inflammation. Collectively, this biology leads to multi-organ damage and premature death in the fifth decade of life. Individuals with SCD often suffer from both acute pain episodes^4–8^ and chronic (steady state) pain^9–13^ that cannot be explained by known SCD pathology^13–16^ such as avascular necrosis or chronic leg ulcers. Opioids are the main analgesic used to treat both acute and chronic SCD pain; however, they are often ineffective and fraught with serious aversive short and long-term side effects^16–19^. SCD disproportionately affects Black or African American individuals who also experience multifactorial health care disparities that are further compounded when seeking pain treatment as racial minorities, especially Black individuals, are less likely to have their pain treated when seeking care^20–22^. These treatment disparities and the current opioid crisis create an immediate need for effective non-opioid based SCD pain therapies. However, in order to develop novel therapies, a better understanding of the neurobiological mechanisms underlying human SCD pain is needed.

Currently, there are humanized mouse models of SCD that exhibit pronounced acute and chronic pain behaviors^23–25^ that include hypoxia/reoxygenation (H/R)-inducible acute pain^26^, spontaneous chronic pain^27–29^, and chronic baseline hypersensitivity to both mechanical and cold stimuli^24,30^. However, growing evidence indicates that there are striking differences in gene expression, transcriptional regulators, and functional responsiveness between human and mouse dorsal root ganglia (DRG) sensory neurons^31–39^. For example, the classic nociceptor capsaicin-sensitive channel TRPV1 is expressed by over twice as many human DRG neurons (75%) as mouse neurons (32%)^36^, and the recently characterized SCD pain target TRPC5 is expressed by significantly more human neurons (75%) than mouse neurons (15%)^37,40^. In contrast, the nociceptor channel TRPA1 is found in far fewer human DRG neurons (16%) than mouse neurons (55%)^36^. To overcome this barrier, human DRG tissues have been suggested as a translational bridge between preclinical rodent models and clinical testing of targets^36,39,41–44^. However, obtaining DRGs from individuals with SCD is virtually impossible and highly impractical due to the relatively low disease incidence and rare clinical indications for DRG resection. Additionally, surgical resections or postmortem tissues offer analysis of static, endpoint molecular signatures and do not allow for longitudinal mechanistic insight into pathogenic processes.

Human induced pluripotent stem cell (iPSC)-derived sensory neurons (iSNs) have recently been used to model chronic pain disorders including chemotherapy-induced peripheral neuropathy^45^. iPSC-derived iSNs express canonical markers of human nociceptors^46–55^ and exhibit functional responses to noxious stimuli similar to native mammalian nociceptors^51,53,55^. Here, we differentiated iSNs from iPSCs derived from individuals with SCD as well as age- and race-matched healthy controls (HCs) to characterize baseline receptor expression and function of SCD iSNs compared to HC iSNs. We also examined SCD patient plasma-mediated effects on SCD iSN hypersensitivity to begin defining extrinsic mechanisms that may sensitize human SCD iPSC nociceptors. Importantly, this study establishes a human-specific model of SCD that proposes mechanisms of SCD pain that may extend beyond a blood-based disease pathology. Moreover, this work allows for the assessment of intrinsic properties of SCD nociceptors that may contribute to chronic baseline sensitivity and provides an unprecedented view of human SCD sensory neuron function.

## Materials and Methods

### Pluripotent stem cells

Three healthy control (HC) iPSC lines (UCSD079i-1-12, PENN062i-278-2, PENN022i-89-1) and three sickle cell disease (SCD) lines (CREM004i-SS2-1, CREM017i-SS19-2, CREM032i-SS48-1) were utilized in these experiments (Table S2). HC lines were age- and race-matched with SCD lines, and all lines were purchased from WiCell. SCD lines were physician diagnosed with HBB(E6V) sickle cell anemia SS according to WiCell. All pluripotent stem cells were maintained on Matrigel (Corning) or Geltrex (Gibco) in Essential 8 (Gibco) and passaged every 4-6 days. iPSCs were karyotypically normal, and differentiated cells were confirmed mycoplasma negative.

### Sensory neuron differentiation

Sensory neurons (iSNs) were differentiated based on the updated Chambers et al. (2012)^53^ protocol provided by Xiong et al. (2021)^45^. Briefly, iPSCs were cultured in E8 on Matrigel coated plates until they reached 80% confluency. Neural differentiation was initiated using KSR medium (80% knockout DMEM, 20% knockout serum replacement, 1X Glutamax, 1X MEM nonessential amino acids, and 0.01 mM β-mercaptoethanol) with dual SMAD inhibition (100nM LDN 193189, 10μM SB 431542). Sensory neurons were patterned using 3μM CHIR99021, 10μM SU5402, and 10μM DAPT starting on day 2 with a stepwise addition of N2 medium (25% DMEM, 25% F12, 50% Neurobasal medium, 1X N2 supplement, 1X B27 supplement) every other day starting on day 4. On day 12, differentiated sensory neurons were dissociated using Accutase (ThermoFisher) and replated onto triple coated (15μg/mL poly-L-ornithine hydrobromide (Sigma), 2μg/mL Laminin (Fisher Scientific), and 2μg/mL fibronectin (Fisher Scientific)) 96-well plates (20,000 cells/well), 6-well plates (480,000 cells/ well), or glass coverslips (52,500 cells/coverslip) in 24-well plates. Differentiated sensory neurons were maintained in N2 media supplemented with neuronal growth factors (10ng/mL human B-NGF, NT3, BDNF, and GDNF). On day 15, cells were treated for 2 hours with freshly prepared mitomycin C (1μg/mL) to eliminate non-neuronal cells. On day 17, medium was completely removed and replaced with fresh medium; subsequent feeds were 50% medium changes every 5-7 days. Sensory neurons were considered mature at day 35 and used for experiments between days 35 and 45.

### Immunocytochemistry

Plated cells were fixed in 4% paraformaldehyde for 20 minutes at room temperature and rinsed with PBS. Nonspecific labeling was blocked and the cells permeabilized with 0.25% Triton X-100 in PBS with 1% BSA and 0.1% Tween 20 for 15 minutes at room temperature. Cells were incubated with primary antibodies overnight at 4°C, then labeled with appropriate fluorescently tagged secondary antibodies. DAPI nuclear dye (Invitrogen) was used to label nuclei. Primary antibodies were guinea pig anti-TRPV1 (ThermoFisher, 1:200), rabbit anti-CGRP (Neuromics, 1:200), and chicken anti-Peripherin (ThermoFisher, 1:500). Secondary antibodies were goat anti-guinea pig Alexa Fluor 488, goat anti-rabbit Alexa Fluor 568, and goat anti-chicken Alexa Fluor 647 (all ThermoFisher, 1:1000 dilutions). Representative images were taken on Zeiss confocal microscope using 20x air objective and are displayed as a maximum intensity projection (MIP) of z-stack image series.

### Quantitative real-time PCR

RNA was isolated from HC and SCD iSN cell pellets using the RNeasy Mini Kit (Qiagen) following manufacturer’s instructions, quantified using a Nanodrop Spectrophotometer, treated with RQ1 Rnase-free Dnase (Promega), and converted to cDNA using the Promega Reverse Transcription system (Promega). SYBR green qRT-PCR was performed in triplicate using cDNA and run on the Bio-Rad CFX384 real time thermocycler. Primers for each target are available in supplemental materials (Table S1). Cq values for each target were normalized to housekeeping gene (GAPDH) values (dCt) and normalized using 1/dCt method. A minimum of three differentiations for each line were collected and run in technical triplicates for each target.

### Calcium imaging

Calcium imaging experiments were performed on HC and SCD iSN following differentiation (days 35-45). Coverslips were incubated in 2.5 μg/mL Fura-2-AM (a dual-wavelength ratiometric calcium indicator dye) in 2% bovine serum albumin for 45 min, followed by a 30 min wash in extracellular normal HEPES buffer (ENH) before imaging. Coverslips were mounted on a Nikon Eclipse TE200 inverted microscope and were superfused with RT ENH (pH 7.4 ± 0.03; 320 ± 3 mOsm; in mM: 150 NaCl • 10 HEPES • 8 glucose • 5.6 KCl • 2 CaCl_2_ • 1 MgCl_2_). Fluorescence images were obtained at 340 nm and 380 nm using Nikon Elements software (Nikon Instruments, Melville, NY). Following a 1 min baseline incubation in ENH, neurons were exposed to 100μM glutamate or 50μM ab-meATP for 2 min, ENH for 3 min, and 50 mM KCl for 1 min. All buffers were superfused at a rate of 6 mL/min. Neurons were included for response magnitude analysis if they displayed a ≥20% increase in the 340/380 nm ratio relative to baseline in response to glutamate, ab-meATP, or KCl.

### Fluo-4 NW calcium flux assay

Calcium flux as an indication of neural activity was measured using the Fluo-4 NW Calcium Assay Kit available through ThermoFisher (#F36206). Assay components were equilibrated to room temperature before 10mL of assay buffer and 100μL of probenecid were added to one bottle of Fluo-4 NW dye mix and vortexed for 1-2min to create the dye loading solution. Growth medium was removed from the 96-well plate using a multichannel pipette, and 100μL of dye loading solution was carefully added to each well. The dye-loaded plates were then protected from light and incubated for 30 minutes at 37°C followed by 30 minutes at room temperature. Just before measuring fluorescence, 25μL of the appropriate agonist or PBS (control) were spiked into each well. Fluorescence (excitation 494nm, emission 516nm) was immediately measured using a GloMax microplate reader with blue filter cube. Relative fluorescence (% above PBS baseline) was calculated by subtracting the fluorescence of PBS-stimulated wells from each test well, then dividing by PBS-stimulated well value and multiplying by 100. Agonist solutions were made fresh each day and included 250mM KCl (final concentration 50mM per well), 500μM L-glutamate (final concentration 100μM per well), 250μM α,β-Methyleneadenosine 5’-triphosphate trisodium salt (ab-meATP, final concentration 50μM per well), and 250μM or 50μM capsaicin (final concentration 50μM or 10μM per well).

### Plasma and recombinant human protein treatments

Human plasma samples were obtained from individuals with SCD during baseline health and during hospitalization for an acute pain event. Plasma was also obtained from healthy Black individuals without SCD. Plasma was collected via routine venipuncture, processed and immediately stored at −80°C. Details of the individuals from whom the plasma samples were obtained are available in supplemental materials (Table S3). Plasma was stored at −80°C and thawed overnight at 4°C on ice before usage; freeze/thaw cycles were limited to 2 for each sample. iSNs were treated with 10% plasma or 100ng/mL recombinant human ET1 (R&D, #1160) or CCL2 (PeproTech, #300-04) diluted in iSN maturation media for 30 minutes at 37°C. After a 30-minute treatment, all media was removed, and 100μL dye loading solution was immediately added to begin the Fluo-4 NW calcium flux assay. Plasma treatments were performed on all lines and variation between lines was nonsignificant (Supplemental Figure S4), so recombinant human protein treatments (ET1 and CCL2) were performed on only PENN022i-89-1 and CREM032i-SS48-1 lines in biological triplicates.

Human samples were collected under an institutional Human Research Review Board approved protocol. Written informed consent was obtained from legal guardians and written informed assent was obtained from the participant when age appropriate.

### Statistical analyses

Experimental conditions within each experiment were performed in technical triplicates for a minimum of three independent experiments unless otherwise noted. Data were analyzed using GraphPad Prism software, and the appropriate statistical tests were performed including the Student’s t-test, 1-way ANOVA, and 2-way ANOVA followed by Tukey’s post hoc analysis of significance. Changes were considered statistically significant when p<0.05.

### Data sharing

For original data, please contact.

## Results

### HC and SCD iSNs express canonical markers of mature and functional human sensory neurons

In order to model SCD, we used three SCD iPSC lines and three age- and race-matched HC iPSC lines. All six lines were capable of generating iSNs following published protocols^45,53^ with few differences in neuronal phenotype (Figure 1A). HC and SCD iSNs expressed similar levels of mRNA transcripts for the mature neuron marker neurofilament heavy chain (NF200) and sensory neuron specific marker calcitonin gene-related peptide (CGRP) as measured by qRT-PCR (Figure 1B). Immunocytochemistry (ICC) confirmed robust protein expression of CGRP, type III intermediate filament protein (Peripherin), and transient receptor potential cation channel subfamily V member 1 (TRPV1) regardless of disease state or line as well as appropriate morphology (Figure 1C, 1D). To further characterize any differences in differentiation efficacy or baseline expression of sensory neuron and pain-relevant ligands and receptors (Table 1) between healthy and disease iSNs, we performed qRT-PCR on the differentiated iSNs. First, we examined the expression of cytokine receptors C-C Motif Chemokine Receptors 1, 2, 3, and 5 (CCR1, CCR2, CCR3, and CCR5) as well as C-C Motif Chemokine Ligand 2 (CCL2) (Table 1A) due to their involvement in peripheral immune response and inflammatory pain^56–60^. We found no significant differences in cytokine receptor transcripts, though we did identify a trend for increased CCL2 transcript production by SCD iSNs compared to HC iSNs (HC=0.0186, SCD=1.0975, p=0.2807).

**Table 1.**
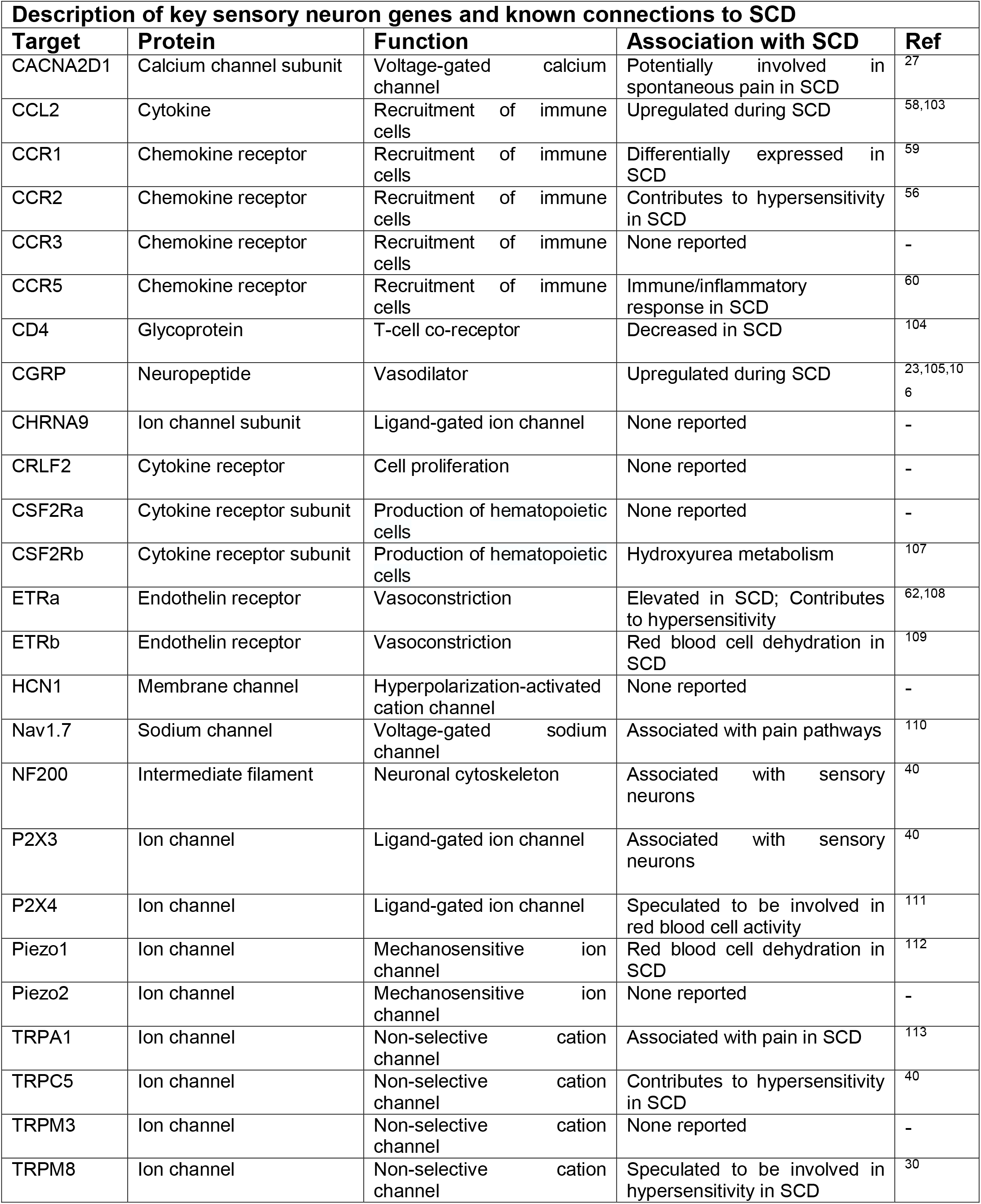
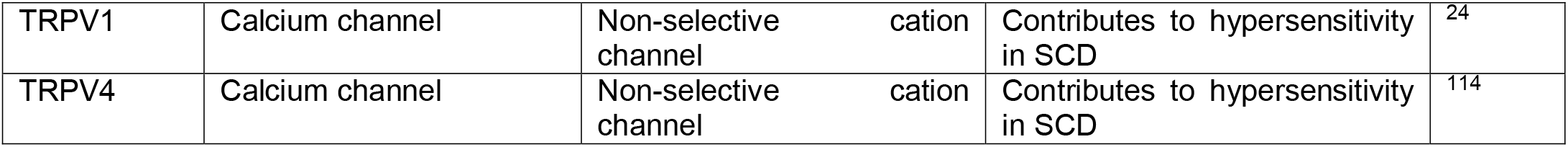
– Description of qRT-PCR targets and any known associations to SCD.

**Figure 1.**
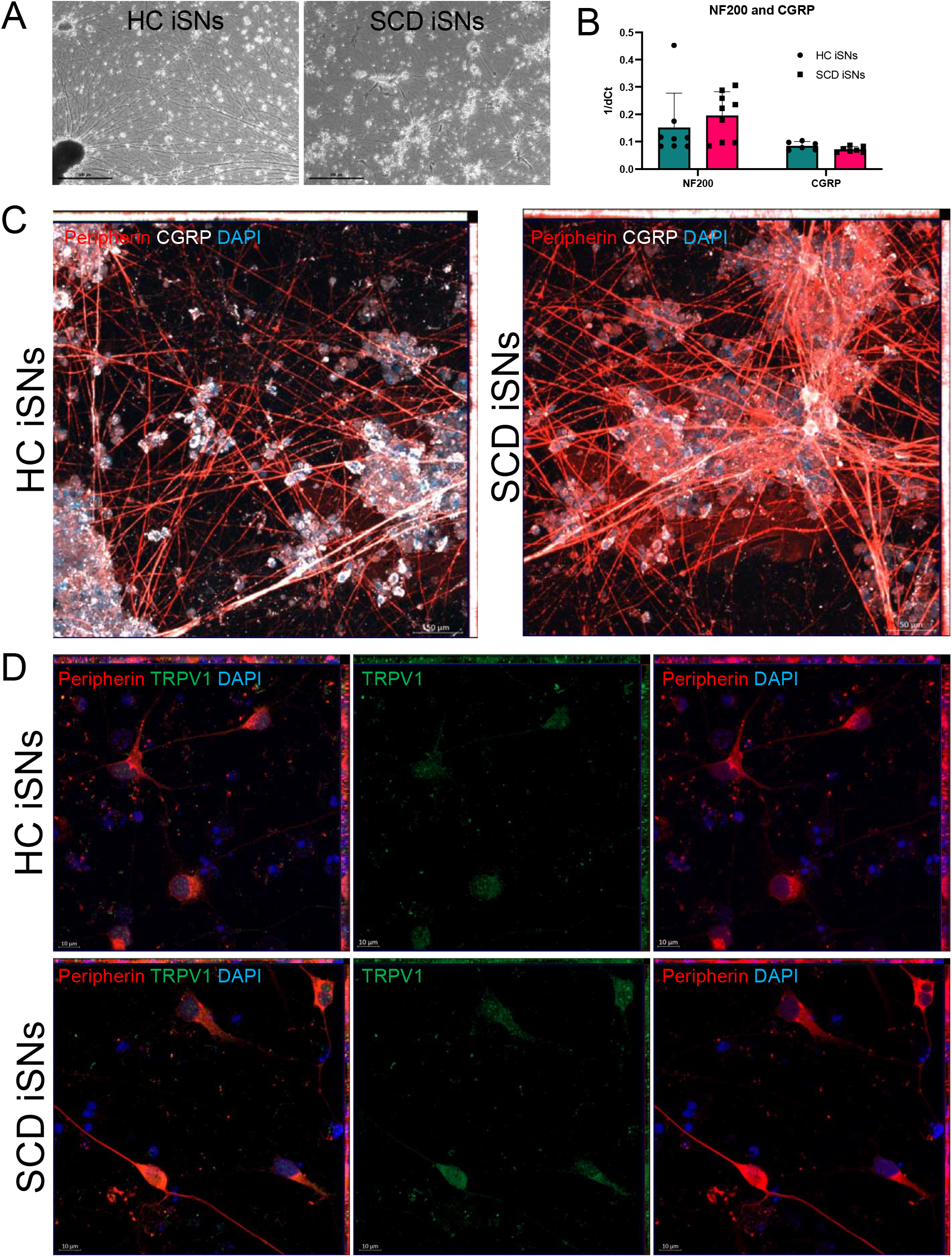
Sickle cell disease (SCD) iPSCs and age-race matched healthy controls (HCs) differentiated into sensory neurons (iSNs) express canonical markers for mature human nociceptors. HC and SCD iSNs (**A**, brightfield images at 4x, scale bar = 500um) express transcripts specific for mature sensory neurons (**B**, NF200 and CGRP) as analyzed via qRT-PCR. **C**. Immunocytochemistry (ICC) shows robust protein expression of Peripherin (red), CGRP (white), and TRPV1 (green) in both HC and SCD iSNs at 20x (**C**, scale bar=50μm) and 63x magnification (**D**, scale bar=10μm). Images are maximum intensity projections (MIPs) from z-stack series taken on Zeiss confocal microscope.

Next, we examined receptors specific for peripheral immune response and vasoconstriction including CD4 molecule (CD4), Cytokine Receptor Like Factor 2 (CRLF2), Colony Stimulating Factor 2 Receptor Subunit Alpha and Beta (CSF2Ra, CSF2Rb), and Endothelin Receptor Type A and B (ETRa, ETRb) (Table 2B). We found little variability in expression between HC and SCD iSN for these transcripts, with the exception of ETRa which showed a trend for higher expression in the SCD iSNs than HCs though with a high degree of variability (HC=0.1256, SCD=0.6931, p=0.5613). Finally, we examined the expression of nociceptor-relevant ion channels including Hyperpolarization Activated Cyclic Nucleotide Gated Potassium Channel 1 (HCN1), Sodium Voltage-Gated Channel Alpha Subunit 9 (Nav1.7), Purinergic Receptor 3 and 4 (P2X3, P2X4), Calcium Voltage-Gated Channel Auxiliary Subunit Alpha2delta 1 (CACNA2D1), and Cholinergic Receptor Nicotinic Alpha 9 Subunit (CHRNA9) (Table 2C). We also examined ion channels suggested to be involved in the SCD pain response including Piezo Type Mechanosensitive Ion Channel Component 1 and 2 (Piezo1, Piezo2), Transient Receptor Potential Cation Channel Subfamily A Member 1 (TRPA1), Transient Receptor Potential Cation Channel Subfamily C Member 5 (TRPC5), Transient Receptor Potential Cation Channel Subfamily M Member 8 (TRPM8), and Transient Receptor Potential Cation Channel Subfamily V Member 1 and 4 (TRPV1, TRPV4) (Table 2D). We found consistent expression of these targets between HC and SCD iSNs. TRPV1 did show a trend for higher transcript expression in SCD iSNs compared to HCs (HC=0.5636, SCD=0.7488, p=0.4368), though the magnitude of this difference was not as striking as those for CCL2 or ETRa. We did investigate the variability between lines and found that it was accounted for by performing biological triplicates for each experiment, with no significant differences between lines (Supplemental Figure S4, p=0.9656). These findings indicate that SCD iPSCs and the age- and race-matched HCs can be consistently differentiated into iSNs expressing canonical and functional markers appropriate to this cell type.

**Table 2.** – Key sensory neuron gene transcript expression analyzed by qRT-PCR for HC and SCD iSNs. No significant differences in transcript expression of cytokine receptors (**A**), immune response and vasoconstriction related receptors (**B**), and ion channels (**C, D**) were identified between HC and SCD iSNs as analyzed by qRT-PCR. Higher trends for differences between HC and SCD iSN gene expression are marked with a diamond (◊, ns). Values for each target are normalized to housekeeping gene (GAPDH) and Ct values for each sample were normalized to a single HC sample value to calculate relative fold change (ΔΔCT) in gene expression. Independent t-tests for each target were used to analyze differences in expression between HC and SCD samples.

| Target | HC iSN expression<br>(mean ± SE) | SCD iSN expression<br>(mean ± SE) | T-test<br>(p-value) |
| --- | --- | --- | --- |
| **A. Cytokines & Receptors** |  |  |  |
| CCR1 | 1.1303 ± 0.8083 | 1.3154 ± 1.0460 | 0.8905 |
| CCR2 | 3.1287 ± 3.1055 | 2.2507 ± 1.9219 | 0.8137 |
| CCR3 | 0.6707 ± 0.5664 | 1.7838 ± 1.7412 | 0.5573 |
| CCR5 | 1.3152 ± 0.8065 | 0.8818 ± 0.4786 | 0.6516 |
| ◇CCL2 | 0.0186 ± 0.5340 | 1.0975 ± 0.6934 | 0.2366 |
| **B. Immune Response & Vasoconstriction** |  |  |  |
| CD4 | 0.7275 ± 0.3015 | 1.1605 ± 0.7263 | 0.5932 |
| CRLF2 | 1.6009 ± 1.1369 | 1.3079 ± 0.7801 | 0.8347 |
| CSF2Ra | 0.4063 ± 0.1305 | 0.7119 ± 0.5496 | 0.6019 |
| CSF2Rb | 0.0385 ± 0.2229 | 0.3847 ± 1.2263 | 0.7878 |
| ◇ETRa | 0.1256 ± 0.6420 | 0.6931 ± 0.6029 | 0.5285 |
| ETRb | 2.1103 ± 1.7300 | 1.8979 ± 1.5764 | 0.9288 |
| **C. Ion Channels – Nociceptor Relevant** |  |  |  |
| HCN1 | 0.7730 ± 0.0928 | 0.7844 ± 0.3887 | 0.9779 |
| Nav1.7 | 0.6744 ± 0.0729 | 0.5603 ± 0.3913 | 0.7813 |
| P2X3 | 1.7744 ± 1.4308 | 1.4527 ± 1.0147 | 0.8570 |
| P2X4 | 0.2109 ± 0.1133 | 0.1079 ± 0.8384 | 0.9059 |
| CACNA2D1 | 0.3918 ± 0.0937 | 0.0371 ± 0.6868 | 0.6222 |
| CHRNA9 | 1.8654 ± 1.3584 | 1.6363 ± 1.0630 | 0.8960 |
| **D. Ion Channels – SCD Pain Specific** |  |  |  |
| Piezo1 | 0.4244 ± 0.6581 | 0.5183 ± 1.0204 | 0.9395 |
| Piezo2 | 0.8643 ± 0.6382 | 0.9337 ± 0.7513 | 0.7360 |
| TRPA1 | 2.7452 ± 2.7365 | 1.6610 ± 1.5499 | 0.7360 |
| TRPC5 | 2.2992 ± 1.5932 | 2.8119 ± 2.5534 | 0.8673 |
| TRPM3 | 1.6454 ± 1.1363 | 1.5149 ± 0.9201 | 0.9301 |
| TRPM8 | 0.0087 ± 0.7276 | 0.0028 ± 0.5776 | 0.9951 |
| ◇TRPV1 | 0.5636 ± 0.0269 | 0.7488 ± 0.2250 | 0.4368 |
| TRPV4 | 0.5997 ± 0.1999 | 0.9690 ± 0.5988 | 0.5718 |

### HC and SCD iSNs are functional and show similar baseline responses to hyperpolarization

We next sought to validate that the differentiation resulted in functional neurons. Both HC and SCD iSNs responded to 50mM KCl, 100μM glutamate, and 50μM α,β-Methyleneadenosine 5’-triphosphate trisodium salt (ab-meATP) using ratiometric calcium imaging analysis (Figure 2A, 2B). No significant differences in percentage of iSNs responding or magnitude of response were found between the HC and SCD iSN in response to these agonists (Chi-squared, 1-way ANOVA, p>0.05). To capture the low percentage of capsaicin responders (expected 2-10%^45^) within the differentiated iSN cultures, we adapted a high-throughput calcium flux assay (Fluo4-NW) to measure total intracellular calcium of 20,000 iSNs within each well of a 96-well plate in response to the agonist treatment. Using this assay, we confirmed that there were no differences in HC and SCD iSN response to 50mM KCl (Figure 2C, p>0.05), 100μM glutamate (Figure 2D, p>0.05), or 50μM ab-meATP (Figure 2E, p>0.05). Both HC and SCD iSNs were found to have consistent responses to 10μM capsaicin, with no differences between HC and SCD iSNs (Figure 2F, p>0.05). We also included a 50μM capsaicin condition (Figure 2G) and found that HC iSNs had a nonsignificant trend for increased calcium flux in response to 50μM capsaicin compared to 10μM, whereas SCD iSNs responded similarly to both concentrations. These data confirm the iSNs respond to stimulation with an increase in intracellular calcium associated with neuronal firing and validate the ability to capture responses in a high-throughput assay.

**Figure 2.**
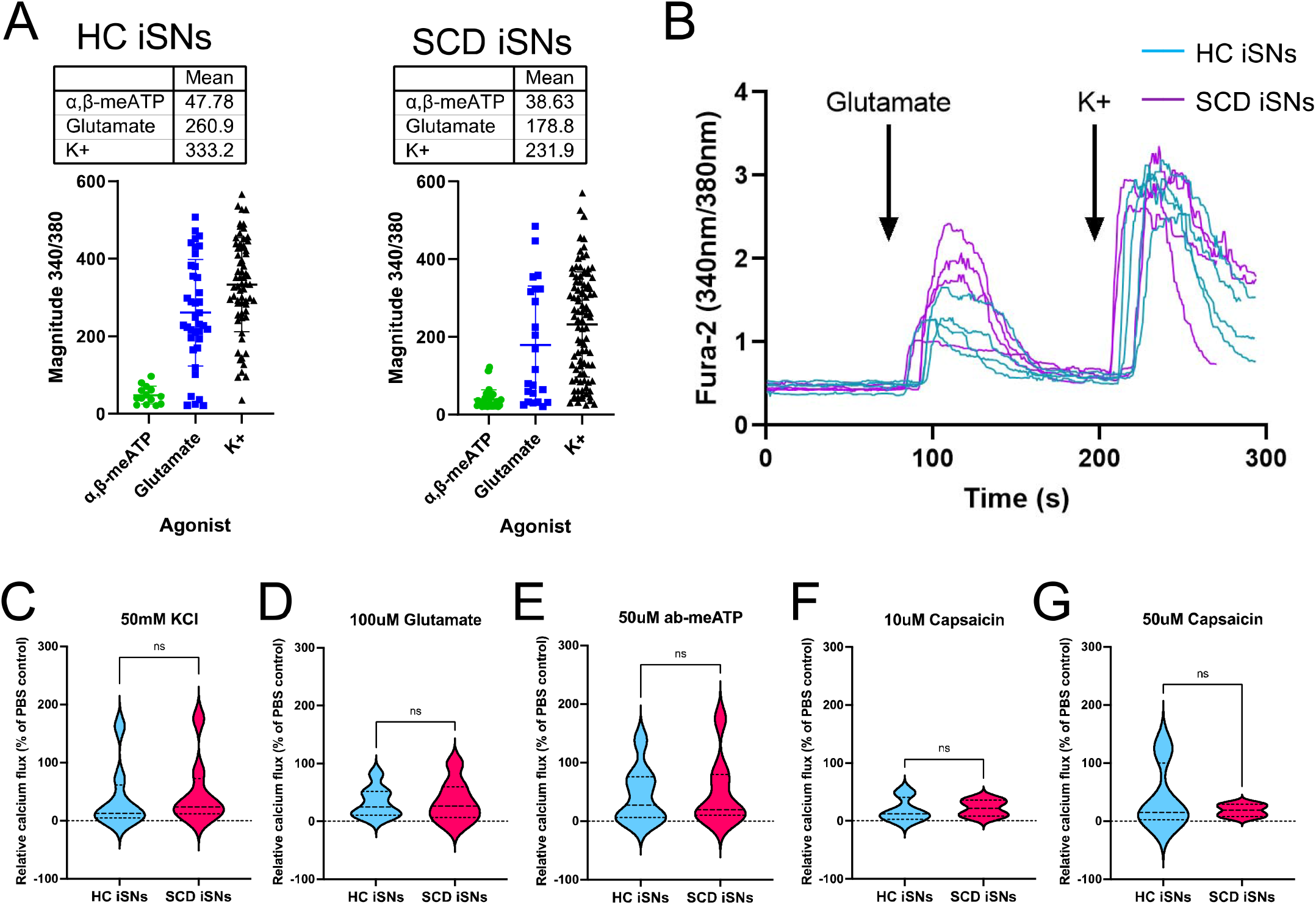
Functional characterization of HC and SCD iSNs using traditional and high-throughput calcium assays. HC and SCD iSNs respond to 50mM KCl, 100μM glutamate, 50μM ab-meATP, and 10μM / 50μM capsaicin in traditional calcium imaging setup (**A, B** representative traces) as well as high-throughput Fluo4-NW calcium flux assay (**C-G**). No significant differences are found between HC and SCD iSN response to these agonist treatments at baseline. Notably, SCD iSNs had similar responses to 10μM and 50μM capsaicin, while HC iSNs show a trend for higher responses to 50μM capsaicin than 10μM. Student’s t-test, ns.

### Human plasma differently sensitizes HC and SCD iSNs

We then investigated the effect of human plasma from individuals with and without SCD on iSN hypersensitivity. In SCD, the blood nerve barrier of DRGs can be compromised and become leaky in chronic pain conditions^61^, thereby allowing circulating ligands access to receptors and ion channels on peripheral neuronal processes. To address this external effect on iSN hypersensitivity, we treated HC and SCD iSNs with plasma taken from healthy individuals without SCD (HC plasma, n=4), individuals with SCD during baseline health (SS BL plasma, n=4), or individuals with SCD during hospitalization for acute pain (SS IP, n=4). Importantly, HC plasma was age- and race-matched to that from individuals with SCD. Overall, we found no significant differences between the plasma treated HC and SCD iSNs in response to 50mM KCl (Figure 3A), 100μM glutamate (Figure 3B), and 50μM ab-meATP (Figure 3C) (2-way ANOVA, p>0.05). Importantly, SCD iSNs did show a significant increase in response to 10μM capsaicin after SS IP plasma treatment compared to HC iSNs (Figure 3D, p=0.0240). This difference was lost in the 50μM capsaicin condition (Figure 3E, p>0.05), suggesting increased sensitivity of SCD iSNs to painful stimuli after exposure to SS IP plasma may be showing maximum response at 10μM capsaicin. When examining the effect of the plasma treatment on responses within HC and SCD iSNs, we found that SS IP plasma increased SCD iSN response to 100μM glutamate (Figure 3F left, 2-way ANOVA, p=0.0435) and 10μM capsaicin (Figure 3F right, p=0.0266) compared to untreated (UTX) SCD iSNs. HC iSNs did not show these same effects but did have increased response to 50μM ab-meATP after SS IP plasma treatment compared to UTX (p=0.0405) and HC plasma treated conditions (p=0.0089). These data reveal possible intrinsic differences within SCD iSNs that may contribute to increased pain sensation during SCD acute pain (Figure 4A).

**Figure 3.**
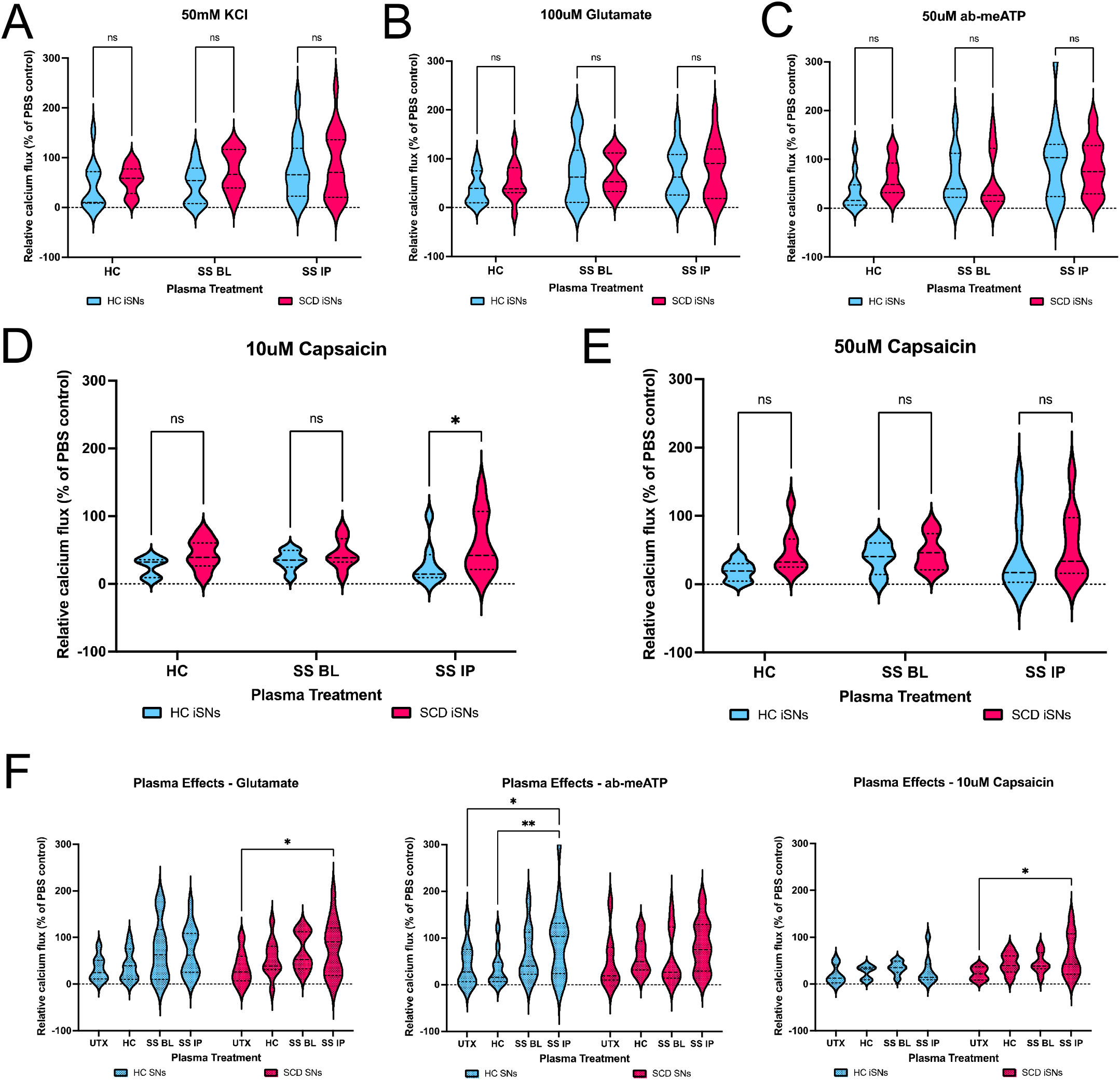
Patient plasma differentially sensitizes HC and SCD iSNs to agonist stimulation. HC and SCD iSNs treated with plasma samples from healthy patients (HC), sickle cell patients at baseline (SS BL), or sickle cell patients experiencing an acute pain crisis (SS IP) show changes in functional response to agonists. No significant differences in response to KCl (**A**), glutamate (**B**), or ab-meATP (**C**) were found between HC and SCD iSNs after any plasma treatments (2-way ANOVA, ns). **D**. Treatment of SCD iSNs with SS IP plasma did significantly increase response to 10μM capsaicin compared to HC iSN response after SS IP plasma treatment (2-way ANOVA, *p<0.05). **E**. No differences found between HC and SCD iSN response to 50μM capsaicin after plasma treatment (2-way ANOVA, ns). **F**. Data from **B-D** represented to compare effects of plasma on iSN response. SS IP plasma significantly increased SCD iSN response to glutamate (**F**, left) and 10μM capsaicin (**F**, right) compared to UTX SCD iSNs, but in HC iSNs significantly increased response to ab-meATP only (**F**, middle). 2-way ANOVA, *p<0.05, **p<0.005.

**Figure 4.**
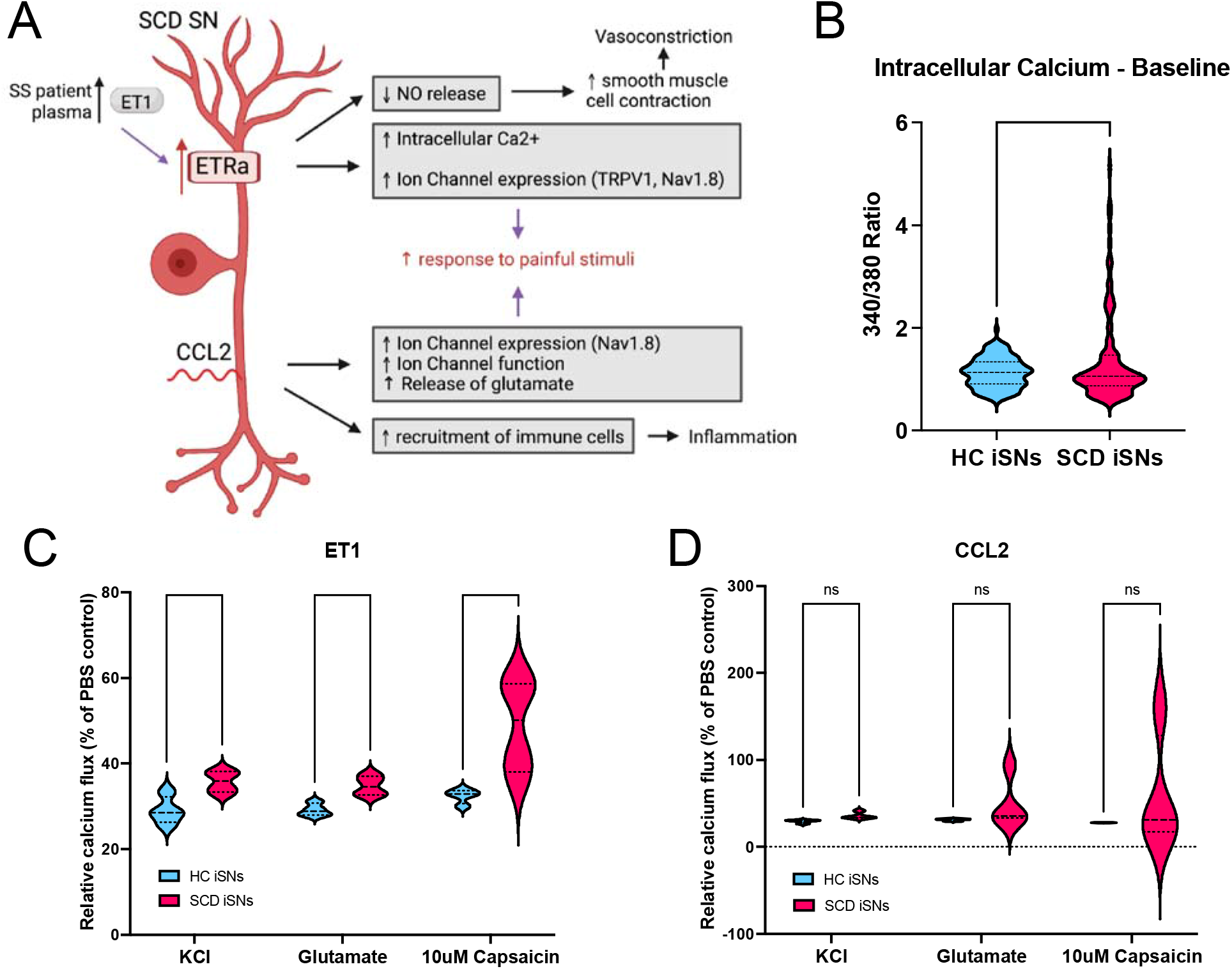
Proposed role of ETRa and CCL2 in SCD iSN hypersensitivity to painful stimuli. **A**. Schematic of novel insight into SCD pathogenesis revealed using human iPSC model and connections to previously published mechanisms. **B**. Post-hoc analyses reveal high degree of variability and abnormal baseline intracellular calcium levels in SCD iSNs compared to HCs (t-test, **p<0.01). Treatment of iSNs with 100ng/mL recombinant human ET1 (**C**) induced significantly higher responses to 50mM KCl, 100μM glutamate, and 10μM capsaicin in SCD iSNs than HC iSNs (t-tests, *p<0.05. Treatment of iSNs with 100ng/mL recombinant human CCL2 (**D**) did not induce this change but did increase variability of SCD iSN responses compared to HC iSNs (t-tests, ns). Key for **A**: findings establishing using this human iPSC model are shown in red, purple arrows are suggested connections to previously published data (gray boxes).

### SCD iSNs express phenotypes consistent with patient populations and mouse models of SCD

We next aimed to further explore potential mechanisms for the increased responses to SS IP plasma. Post-hoc analyses of our calcium imaging data revealed variable calcium response levels and a significant number of SCD iSNs with abnormally high baseline calcium levels compared to HC iSNs (Figure 4B, p=0.0024) suggesting that SCD iSNs are residing in an already sensitized state Previous studies have also shown that plasma from individuals with SCD contains increased levels of endothelin 1 (ET1)^62–67^. Signaling of ET1 through increased ETRa receptors in SCD iSNs can contribute to decreased nitric oxide release^68,69^ contributing to vasoconstriction^70,71^, increased intracellular calcium^71,72^, and increased expression of ion channels like TRPV1^73,74^ and Sodium Voltage-Gated Channel Alpha Subunit 10 (Nav1.8)^62,75^. Moreover, increased CCL2 production by SCD iSNs can also result in increased neuronal excitability^76,77^, ion channel expression (Nav1.8) and function^78^, release of glutamate^79^, and recruitment of immune cells resulting in inflammation^57,80^. Therefore, we added 100ng/mL of recombinant human ET1 (Figure 4C) and CCL2 (Figure 4D) into HC and SCD iSN cultures. ET1 significantly increased calcium flux of the SCD iSNs in response to 50mM KCl (p=0.0333), 100μM glutamate (p=0.0186), and 10μM capsaicin (p=0.0333) compared to HC iSNs. CCL2 did not induce these direct increases in calcium flux (p>0.05), though it did increase the range of calcium responses of SCD iSN to glutamate and 10μM capsaicin. These data support the idea that increased expression of certain receptors and ligands in SCD iSNs can contribute to increased sensitivity to circulating plasma factors during acute SCD pain.

## Discussion

This study presents SCD patient-derived iPSCs differentiated into iSNs as a model system for studying intrinsic and dynamic mechanisms of nociception in SCD. In our initial characterizations, we established that HC and SCD iSNs look and function like human nociceptors (Figures 1, 2). Although we did not identify any significant differences in markers of nociceptors or targets relevant to SCD pain between HC and SCD iSNs (Tables 1, 2), we did note that SCD iSNs have increased and highly variable baseline calcium levels (Figure 4B) compared to HC iSNs. This may indicate populations of SCD iSNs that are sensitized at baseline and exhibit increased spontaneous activity^81^.

Further, we found the ability of SCD patient plasma to sensitize iSNs (Figure 3). Untreated SCD iSNs appear to reach a maximum calcium response to capsaicin at 10μM, whereas HC iSNs have a greater calcium response to 50μM than 10μM capsaicin (Figure 2F, 2G). The high calcium baselines seen in the SCD iSNs (Figure 4B) may contribute to this finding, as intracellular calcium has been previously proposed to desensitize sensory neurons to capsaicin^82^. However, when treated with SCD patient plasma, SCD iSNs surpass the untreated maximum in both the 10μM and 50μM capsaicin conditions (Figure 3D, 3E). This suggests a role of plasma factors in overcoming the desensitization of untreated SCD iSNs to TRPV1 agonism, perhaps through localization or sensitization of TRPV1 channels at the plasma membrane^83,84^. Increased ET1^73,74^ and factors associated with hypoxia (H_2_O_2_, TNFα, IL6)^85,86^, hemolysis (hemin)^87,88^, and acidosis (H^+^)^89,90^ are proposed sensitizers of TRPV1 and are also increased during acute SCD pain^91–93^.

Importantly, treatment with SCD patient plasma revealed only one significant difference based on iSN disease state. This was a significantly increased calcium response of the SCD iSNs after treatment with SCD patient plasma collected during an acute pain crisis, and the effect was capsaicin-specific (Figure 3D). The effect is lost with 50μM capsaicin, likely because both HC and SCD iSNs are reaching their maximum response levels (Figure 3E). This finding expands on the prior reports on TRPV1 sensitization in SCD mice^24^ and can be further investigated using molecular studies of iSNs including patch-clamp electrophysiology^94^. TRPV1 localization can also be further studied using channel-tagged reporter proteins^95–97^ to determine the contribution of receptor trafficking on SCD iSN response to capsaicin.

It is also notable that the SCD patient plasma did have effects on the HC iSNs. This implicates factors in SCD patient plasma that are capable of sensitizing healthy nociceptors. For example, although no differences between HC and SCD iSN response to glutamate were found after plasma treatments (Figure 3B), SS IP plasma did significantly increase SCD iSN response to glutamate compared to the untreated SCD iSNs (Figure 3F). A similar phenomenon was found where SS IP plasma significantly increased HC iSN response to ab-meATP compared to untreated and HC plasma treated HC iSNs (Figure 3F), but again this impact was not significantly different between HC and SCD iSNs (Figure 3C). There are several possibilities to explain this nuanced effect. First, if the trend for higher ETRa expression in SCD iSNs is biologically relevant compared to HC iSNs despite not reaching statistical significance, the high levels of ET1 in patient plasma^62–67^ may be key to this relationship. In support of this, treatment with recombinant ET1 significantly increased SCD iSN response to all tested agonists compared to HC iSNs (Figure 4C).

However, ET1 alone does not appear to explain the effects of plasma on SCD iSNs because the increased activity of SCD iSNs after ET1 treatment is not capsaicin-specific (Figure 4C). High dosage of ET1 could contribute to the non-specific hypersensitization, or it may be that a specific combination of plasma factors is required to recreate the capsaicin-specific effect of plasma treatment in SCD iSNs. This hypothesis is supported by the increased variability of SCD iSNs response seen after CCL2 treatment compared to HC iSN response (Figure 4D). Increased CCL2 is associated with glutamate signaling in sensory neurons^79^ and could contribute to the increased response of SCD iSNs to glutamate after treatment SS IP plasma. The ab-meATP-specific effect seen in the HC iSNs after plasma treatment may be due to increased levels of lysophosphatidic acid (LPA) released during vaso-occlusive crises^98^ sensitizing the ATP-activated channel P2X3^99,100^.

An unanswered question remains as to how a genetic mutation in hemoglobin (Hb) could impact iSN function. There is evidence linking Hb expression in neurons to expression of genes associated with mitochondrial function^102^, which provides a precedent for Hb-dependent transcriptional regulation in neurons. Therefore, we can assess Hb expression in the iSNs to investigate its connection to transcriptional changes relevant to sensory neuron function and malfunction.

In summary, we present SCD iSNs as a large-scale and higher-throughput platform to study human-specific SCD nociception in a manner not previously achievable. We have shown that this model recapitulates important disease phenotypes and can be used to investigate individual factors in SCD patient plasma that can contribute to SCD pain sensation and overall neuropathology. We specifically identified ET1 and CCL2 as potential contributors to human nociceptor TRPV1 sensitization in SCD. These data and the iSN system have powerful implications on SCD research. Future studies will leverage the iPSC iSN system described here to identify candidate signaling factors and/or plasma metabolites as mediators of SCD iSN sensitization as well as validate pharmacological and gene therapy approaches to aid therapeutic development.

## Acknowledgements

We would like to thank and acknowledge individuals with and without SCD who contributed to this study. This project was supported by the funding from the National Institutes of Health R01NS070711 (CLS), R37NS108278 (CLS), 1K23HL114636-01A1(AMB), 1R01HL142657-01 (AMB), and Advancing a Healthier Wisconsin (AHW) Endowment (ADE, CLS, AMB).

## Author Contributions

RLA and AB performed and analyzed experiments. All authors designed experiments and interpreted data. AMB, CLS, and ADE supervised the study and provided funding. RLA wrote the manuscript. RLA, AB, and DNT created figures. All authors edited and approved the manuscript.

