## Supplemental tables and figure for "Sickle cell disease patient plasma sensitizes iPSC-derived sensory neurons from sickle cell disease patients"

Supplemental Table S1 – qRT-PCR primer sequences

| **Target** | **Forward Sequence** | **Reverse Sequence** |
| --- | --- | --- |
| NF200 | CTGAGGAACACCAAGTGGGAGA | TCCGACACTCTTCACCTTCCAG |
| CGRP | CTCTAAGGGTGAGCTGAAGCAG | ACCTGTTCATCTTGATTAGCATCC |
| CCR1 | CAACTCCGTGCCAGAAGGTGAA | GTTCAGGAGGTAGATGCTGGTC |
| CCR2 | CAGGTGACAGAGACTCTTGGGA | GGCAATCCTACAGCCAAGAGCT |
| CCR3 | TACTCCCTGGTGTTCACTGTGG | ACGAGGAAGAGCAGGTCCGAAA |
| CCR5 | TCTCTTCTGGGCTCCCTACAAC | CCAAGAGTCTCTGTCACCTGCA |
| CCL2 | CAGCCAGATGCAATCAATGCC | TGGAATCCTGAACCCACTTCT |
| CD4 | CCTCCTGCTTTTCATTGGGCTAG | TGAGGACACTGGCAGGTCTTCT |
| CRLF2 | AAGCGACTGGTCAGAGGTGACA | GAGGAGAGACACCATCAGAAGG |
| CSF2Ra | CCTGTCAGGATTAACGTCTCGC | CATTGCTGGGAGGGTTGAATCG |
| CSF2Rb | TCTTCCTCACCATCGCTGTGCT | CTGGAACAGGTGGCTCTTGCTG |
| ETRa | GCGATTGGCTTCGTCATGGTAC | GAAGATCGCAGTGCACACCAAG |
| ETRb | CAGAAAGCCTCCGTGGGAATCA | ACAGCCAGAACCACAGAGACCA |
| HCN1 | ACGAGAAGGAGCCGTGGGTAAA | ACGTCCTTTGGTCAGCAGGCAA |
| Nav1.7 | GTGGAAGGATTGTCAGTTCTGCG | GCCAACACTAAGGTGAGGTTACC |
| P2X3 | GCGTTTCTGAGAAAAGCAGCGTG | CGGATGCCAAAAGCCTTCAGGA |
| P2X4 | GTGGCGGATTATGTGATACCAGC | CACACAGTGGTCGCATCTGGAA |
| CACNA2D1 | GCTATTCACGGATGGAGGAGAAG | CCATCCACTGAATAGGTCCTCTG |
| CHRNA9 | CTAATGCTCTTCGTCCAGTGGAA | GTGAGATAGGCATCGTGCCAGA |
| Piezo1 | CCTGGAGAAGACTGACGGCTAC | ATGCTCCTTGGATGGTGAGTCC |
| Piezo2 | GACGGACACAACTTTGAGCCTG | CTGGCTTTGTTGGGCACTCATTG |
| TRPA1 | CTCCTCTCCACATAGCTGTGCA | GTGCACGCAATGATCACAGCTGT |
| TRPC5 | CCACCAGCTATCAGATAAGG | CGAAACAAGCCACTTATACC |
| TRPM3 | GGAAGTGTTTGCGGACCAGATAG | GCACGATCCAAGCTCCTGTCTT |
| TRPM8 | CTGGTTGCGAACTTCCGAAGAG | GGTGCCGAGTAATAGGAGACAC |
| TRPV1 | ACACCTGATGGCAAGGACGACTAC | AAAAGGGGGACCAGGGCAAAGTT |
| TRPV4 | TCACTCTCACCGCCTACTACCA | CCCAGTGAAGAGCGTAATGACC |

Supplemental Table S2 – iPSC line donor demographic information

| **iPSC line** | **Disease** | **Genetic Alteration/ Mutation** | **Age** | **Gender** | **Ethnicity** |
| --- | --- | --- | --- | --- | --- |
| CREM004i-SS2-1 | Sickle cell anemia SS | HBB(E6V), A>T mutation in beta globin gene | 32 | F | African American |
| CREM017i-SS19-2 | Sickle cell anemia SS | HBB(E6V), A>T mutation in beta globin gene | 30 | M | African American |
| CREM032i-SS48-1 | Sickle cell anemia SS | HBB(E6V), A>T mutation in beta globin gene | 30 | M | African American |
| UCSD079i-1-12 | None reported | N/A | 32 | F | African American, Latino |
| PENN062i-278-2 | None reported | N/A | 33 | M | African American |
| PENN022i-89-1 | None reported | N/A | 28 | M | African American |

Supplemental Table S3 – Patient sample demographic information

|  | **Age** | **Sex** | **Race** | **Genotype** |
| --- | --- | --- | --- | --- |
| **Healthy Controls** | | | | |
| HC 1 | 17 | F | Black/African American | N/A |
| HC 2 | 11 | M | Black/African American | N/A |
| HC 3 | 14 | F | Black/African American | N/A |
| HC 4 | 13 | F | Black/African American | N/A |
| Mean age = 13.75 | | | | |
| **Individuals with Sickle Cell Disease** | | | | |
| Baseline state of health | | | | |
| SCD 2 BL | 15 | F | Black/African American | SS |
| SCD 3 BL | 11 | M | Black/African American | SS |
| SCD 4 BL | 11 | M | Black/African American | SS |
| Mean age = 12.33 | | | | |
| Acute pain (inpatient) | | | | |
| SCD 1 IP | 16 | F | Black/African American | SS |
| SCD 2 IP | 16 | F | Black/African American | SS |
| SCD 3 IP | 11 | M | Black/African American | SS |
| SCD 4 IP | 9 | M | Black/African American | SS |
| Mean age = 13 | | | | |

Supplemental Figure S4 – Calculated variability between iPSC lines and biological replicates

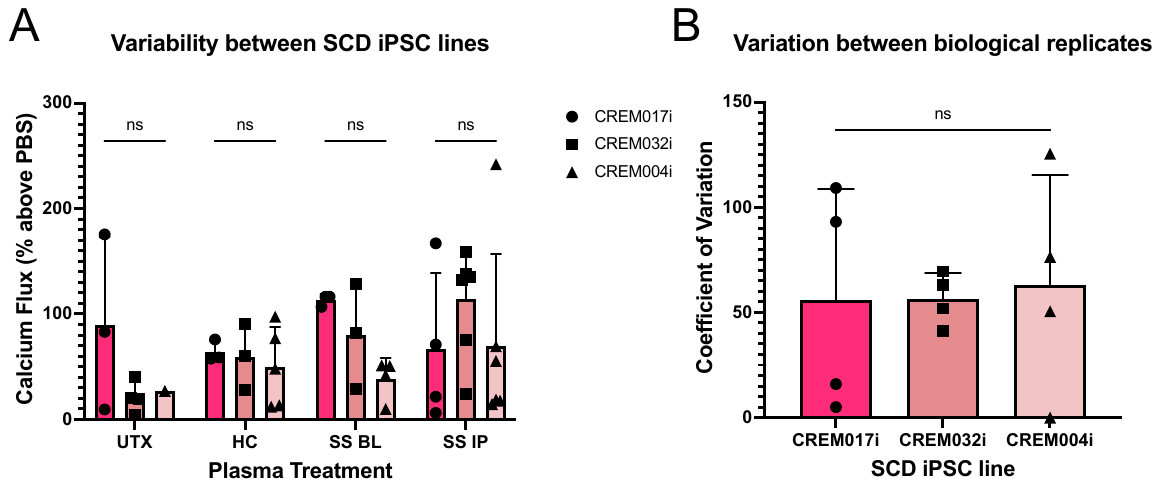

**Figure S4.** **SCD patient-derived iPSCs differentiated into iSNs display nonsignificant variability between independent lines.** **A**. No significant differences in the calcium flux responses of 3 independent SCD iPSC lines differentiated into iSNs after 50mM KCl stimulation (2-way ANOVA, variation due to iPSC line = 5.72%). **B**. Residual variability between lines likely due to differences between biological replicates, with no significant differences found between biological replicate variability between lines (1-way ANOVA, p=0.9656). All experiments performed in biological triplicates to account for this variability.
